# Gamma oscillations across recording scales show a preference for saturated long-wavelength (reddish) hues in the primate visual cortex

**DOI:** 10.64898/2026.01.22.701022

**Authors:** Ankan Biswas, Supratim Ray

## Abstract

Stimulus-induced narrowband gamma oscillations (30–70 Hz) arise due to excitatory-inhibitory interactions and have been proposed to carry feedforward prediction errors during visual processing. However, the dependence of gamma on stimulus color is not well characterized, with some studies showing a preference for saturated long-wavelength (reddish) hues in invasive recordings from the primary visual cortex (V1), while others showing no preference for red in luminance and cone-contrast balanced colors spanning the Derrington-Krauskopf-Lennie (DKL) isoluminant plane in non-invasive magnetoencephalography (MEG) recordings. To address these discrepancies, we simultaneously recorded local field potentials (LFPs) from V1 (n=2 monkeys) along with scalp electroencephalography (EEG), while presenting luminance-matched hues spanning the entire permissible range of colors. We found that saturated reddish hues produced the strongest gamma in both LFP and EEG in both monkeys. However, gamma was reduced when the colors were desaturated, so that selectivity for reddish hues was diminished or absent on the DKL space, as shown previously. Interestingly, selectivity was further reduced in EEG compared to LFP. Simultaneous recordings from an intermediate visual area (V4, n=1) revealed weak color-induced gamma, partially explaining the weakening of hue selectivity in macro-signals. Gamma power increased with increasing activation along the red-green (L-M) cardinal axis but was virtually absent along the blue-yellow (S-(L+M)) axis, suggesting that gamma could be a reflection of the L-M cone-contrast mechanism. These results comprehensively resolve the earlier discrepancy and shed light about the neural mechanisms underlying gamma generation in primate V1.

**Significance Statement:** Gamma-band brain rhythms are markers of cortical computation and disease, yet their dependence on color has remained controversial because studies using different recording methods and stimulus color spaces yielded conflicting results. By densely sampling colors across the entire monitor gamut in an isoluminant space and mapping brain rhythms across multiple neural scales in primates, we show that primary visual cortex generates exceptionally strong gamma for saturated reddish hues. Crucially, this effect is driven by contrast in specific cone pathways (specifically, L-M cone contrast), and the selectivity diminishes in desaturated color spaces, downstream visual areas and scalp recordings. This explains why the “red bias” observed invasively often vanishes in non-invasive recordings and demonstrates that color-induced gamma reflects pathway-specific circuit engagement.

## Introduction

Narrowband gamma oscillations (30–70 Hz) in visual cortex arise from recurrent interactions between excitatory neurons and inhibitory interneurons (1–3). Since these oscillations provide a functional readout of cortical excitation-inhibition (E–I) balance, they have emerged as critical targets for elucidating sensory computation mechanisms and identifying biomarkers for neurodegenerative pathologies, including aging and Alzheimer’s Disease (4–8). Further, recent studies have hypothesized that gamma oscillations could carry feedforward prediction-error signals in a predictive coding (9–12) or predictive routing scheme (13, 14). To test these predictions and to better understand the circuit mechanisms behind gamma generation, it is important to understand the dependence of gamma on features of natural images (15–19).

In the primary visual cortex (V1), gamma response mapping has been thoroughly done using achromatic gratings, and found to be dependent on a variety of features such as size, contrast, spatial frequency, orientation, and temporal frequency (20–27). Gamma responses to chromatic stimuli are less well studied. Some studies using microelectrodes showed that large, highly saturated long-wavelength (reddish) hues can induce unusually high gamma responses that were much stronger than those induced by other hues or achromatic stimuli (16, 28, 29). However, these saturated colors were not matched in luminance and were not equated for the response they evoked in the lateral geniculate nucleus (LGN) that provides input to V1. Neural responses in early visual pathways scale with cone contrast—the relative modulation of L, M, and S cone excitations away from the adapted background—rather than absolute wavelength or saturation (30, 31). To control for these factors, Derrington, Krauskopf, and Lennie (1984) developed a color space (DKL) whose axes correspond to the response properties of neurons in the LGN: an achromatic “luminance” axis (L+M), a “red-green” chromatic axis (L−M), and a “blue-yellow” chromatic axis (S−(L+M)). Modulation along each cardinal axis selectively activates one mechanism while leaving the others unchanged, allowing chromatic mechanisms to be isolated independently. A recent study showed that in Magnetoencephalogram (MEG), preference for reddish hues is absent when hue stimuli are taken from the equiluminant DKL space (i.e., are equated for cone-contrast; (32)).

Although these results extend our understanding of the color dependence of gamma, there are two caveats. First, DKL stimuli are constrained to low saturation levels and span a relatively small range of colors that can be rendered on a screen. Since gamma is known to reduce substantially once colors are desaturated (28), the selectivity for reddish hues could be reduced in a desaturated color space. Second, while the preference for reddish hues was originally shown in invasive microelectrode or electrocorticogram (ECoG) recordings in V1, the studies showing a lack of selectivity used MEG, where higher spatial summation of signals from different visual areas could have diminished the selectivity observed in V1.

To resolve this, we performed simultaneous electrophysiology in two macaque monkeys across different neural scales. We recorded local field potentials from V1 in both monkeys and V4 in one monkey. Concurrent scalp EEG was also acquired from both monkeys. We presented isoluminant chromatic stimuli using four different experiments: radial saturation modulation toward monitor primaries, hues spanning the DKL isoluminant ellipse, systematic variation along the L–M cardinal axis, and dense sampling across the accessible CIE color gamut. By combining these experiments, we were able to isolate the contribution of cone-opponent signals to gamma oscillations and characterize how chromatic selectivity transforms across neural scales.

## Results

Since strong gamma oscillations were observed in a previous report (28) using a saturated red hue, we first investigated how gamma oscillations depend on color saturation by varying the stimuli radially from the achromatic white point toward each of the three monitor primaries (red, green, and blue) on an isoluminant plane. We used a low luminance (mean luminance 8.17 ± 0.30 cd/m²) for all the colors (including the pre-stimulus background gray) to allow blue to reach full saturation within the monitor gamut (Figure 1A) as the maximum achievable luminance for pure blue on our display was 8.79 cd/m^2^. Next, we sampled 36 isoluminant colors uniformly around the largest DKL ellipse that we were able to render on our monitor (see Methods and Appendices for more details), maintaining luminance at ∼59.45 cd/m² (Figure 1B). In the third experiment, we varied L–M cone contrast systematically along the cardinal red–green axis under isoluminant conditions (∼60.27 cd/m²), directly testing whether gamma scales with cone-opponent signals (Figure 1C). Finally, we conducted a dense sampling across the accessible color space at low background luminance to comprehensively map gamma responses throughout the full range of hues and saturations (Figure 1D).

**Figure 1:**
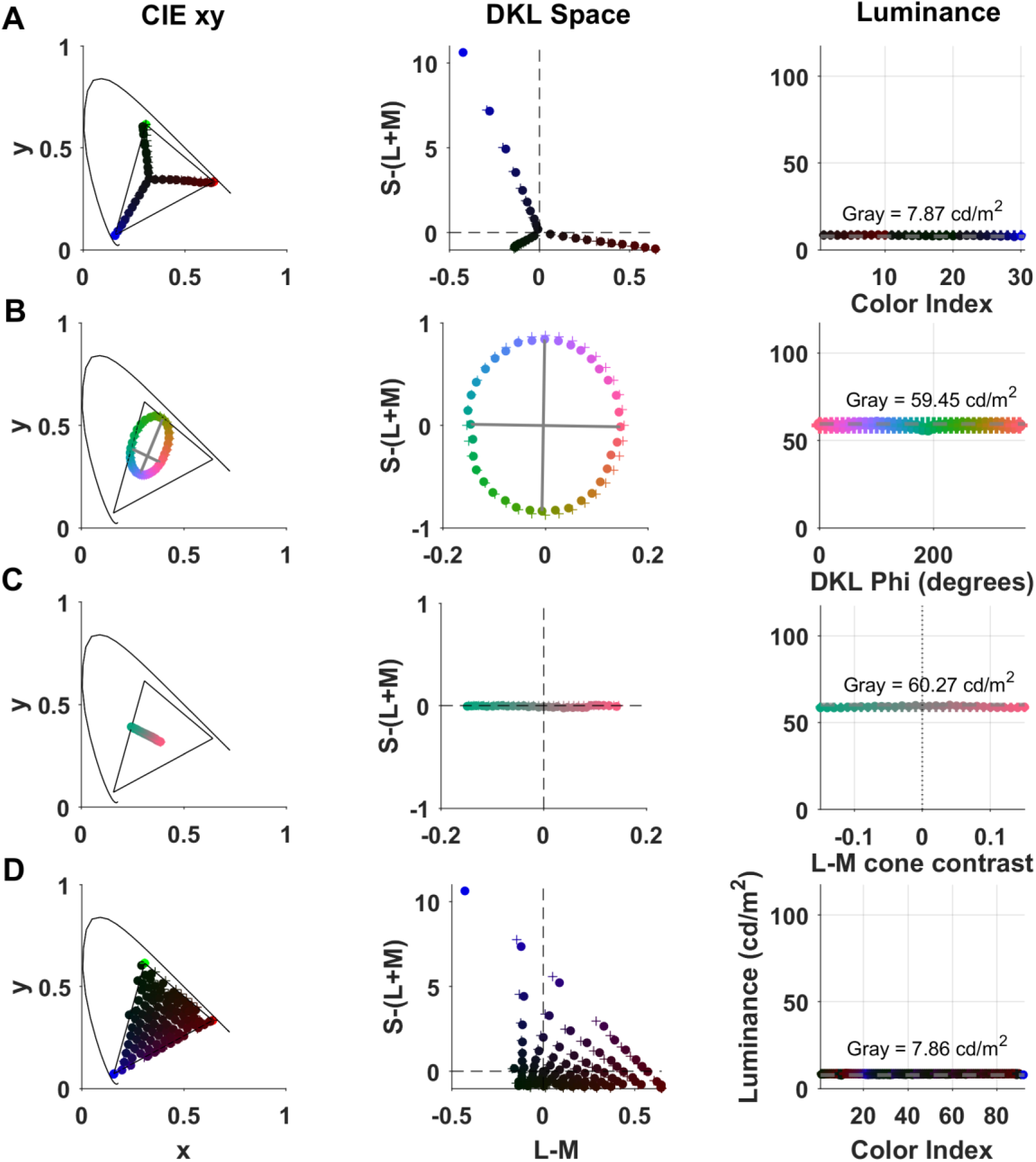
Stimuli in CIE and DKL color spaces along with luminance. Stimuli for four experiments (A–D, rows) shown in CIE 1931 xy chromaticity space (left column), DKL isoluminant plane with L-M and S-(L+M) axes (middle column) and measured photometric luminance (right column). Black curves in CIE space indicate monitor gamut. Colors in DKL space indicate azimuth angle. Desired and actual (measured using a spectroradiometer) colors are shown using “+” and ‘o’ markers, which are largely overlapping. a) Radial saturation variation: stimuli varying radially from the achromatic white point toward the three monitor primaries (red, green, blue) at low background luminance (7.87 cd/m²) where blue reaches full saturation. Middle panel shows these colors in the DKL space; left shows them in CIE xy coordinates. b) Isoluminant hue ellipse: uniform sampling of 36 DKL azimuth angles. Measured luminance is constant across all hues (right; background = 59.45 cd/m²). c) L-M cone contrast variation: stimuli varying systematically along the L-M cardinal axis at multiple cone contrast levels. Luminance remains constant across all contrast levels (right; background = 60.27 cd/m²). d) Dense color space sampling: finer radial-angular sampling across the entire accessible color space at low background luminance (7.86 cd/m²).

### Saturated red produces stronger gamma than other primaries in both LFP and EEG

We presented full-field stimuli at ten incremental saturation levels along each primary color direction (Figure 2, top panel). Gamma oscillations became stronger with increasing saturation for all colors (as shown previously in Figure 4 of Ref (28)), although the colors were not isoluminant in that case), as shown in averaged time-frequency change in power spectra for LFP recordings from V1 (rows 1 and 2; n=31 and 39 sites for M1 and M2) and EEG (rows 3-4, n=6 electrodes for both M1 and M2; Supplementary Figure 1, Electrodes, M1: 1, 2, 4, 6, 8, 10; M2: 1, 2, 3, 5, 8, 10). More importantly, at the highest saturation, red evoked much stronger narrowband gamma oscillations as compared to green or blue. We also observed an increase in centre frequency with increasing saturation. This was observed in both monkeys and for both LFP and EEG recordings.

**Figure 2:**
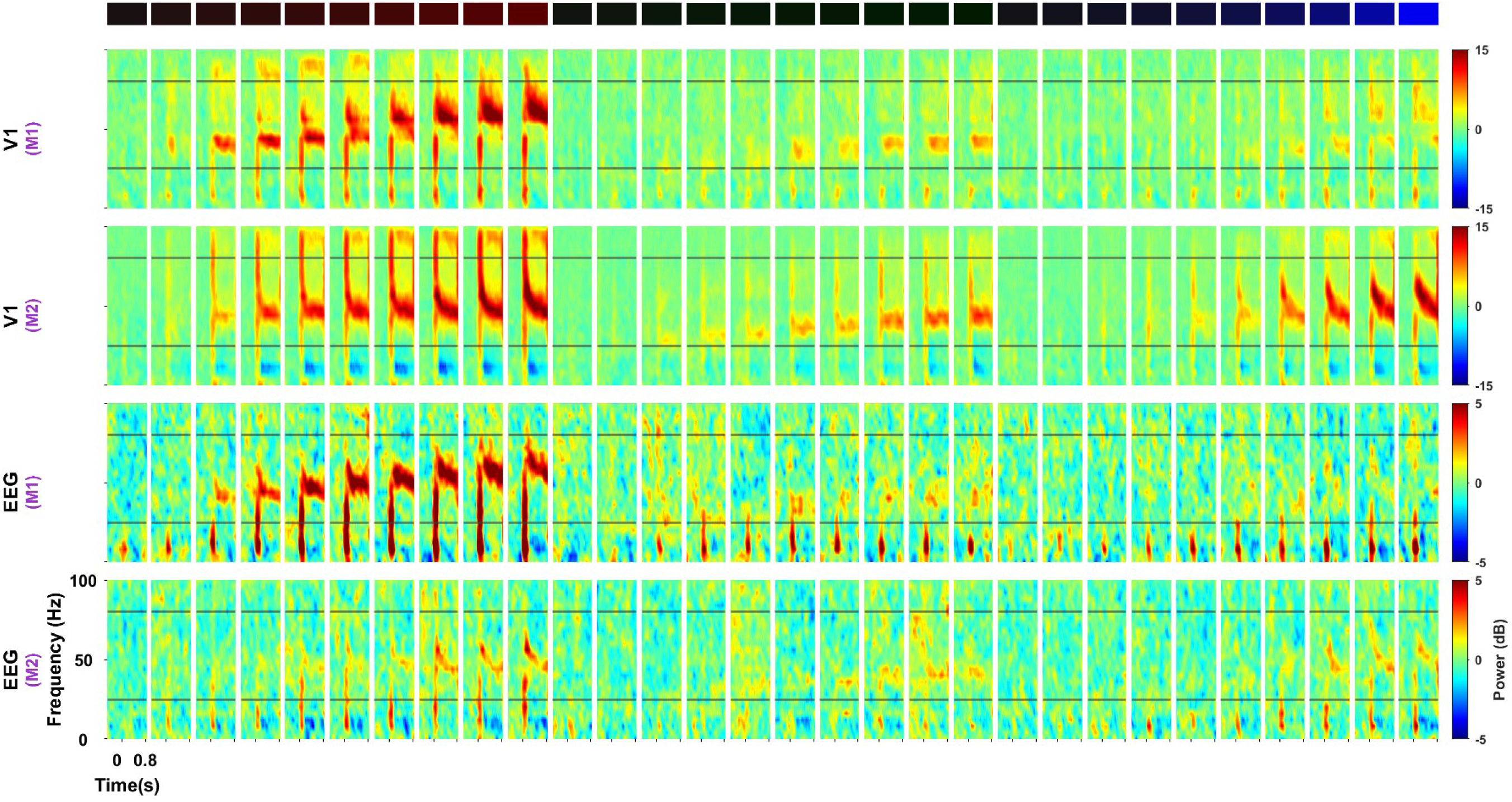
Gamma oscillations during radial saturation variation in CIE color space. Time–frequency spectra showing the decibel change in power from the baseline period (−0.5 s to 0 s, where 0 indicates stimulus onset) for stimuli varying radially from the achromatic white point toward the three monitor primaries (red, green, blue) at 10 saturation levels each (indicated by color swatches at top, progressing from white to saturated red, green, and blue). Rows show population averages from V1 LFP (rows 1–2; n = 31 and 39 sites for Monkey 1 (M1) and Monkey 2 (M2), respectively) and scalp EEG (rows 3–4; electrode configuration as described in Methods) recordings from M1 (n=6) and M2 (n=6). Each small panel corresponds to one saturation level along one primary direction. Time axis spans -0.5–0.8 s; frequency axis, 0–100 Hz. Black horizontal lines indicate the gamma band limits. Background luminance was 7.87 cd/m² where the blue primary reaches full saturation.

Population tuning curves —constructed by computing peak gamma power at each saturation level and averaging across all electrodes—confirmed that gamma power increased with increasing saturation for all three primary colors (Figure 3). In both monkeys, peak gamma power increased with saturation for all three colors, with red consistently producing the highest power. The relative ordering of green and blue varied across animals—M1 showed preference for green over blue, while M2 showed the opposite pattern (Figure 3A, B). Statistical analysis at the highest saturation levels (average of levels 8–10 for each color; figure 3C) revealed that red stimuli generated significantly stronger V1 gamma than green (two-tailed paired t-test, M1: t(30)=157.04, p=2.68×10^-45^; M2: t(38)=59.98, p=3×10^-39^) or blue (two-tailed t-test, M1: t(30)=177.70, p=6.61×10^-47^; M2: t(38)=29.34, p=1.03×10^-27^). EEG effects were weaker than V1 but showed significant red–green and red–blue differences in M1 (red–green: *t*(5)=5.23, *p*=3.40×10^−3^; red–blue: *t*(5)=4.34, *p*=7.40×10^−3^), whereas in M2 only the red–blue contrast reached significance (red–blue: *t*(5)=3.19, *p*=0.02; red–green: *t*(5)=1.60, *p*=0.17). The significance in EEG data should be interpreted with caution because of the low sample size.

**Figure 3:**
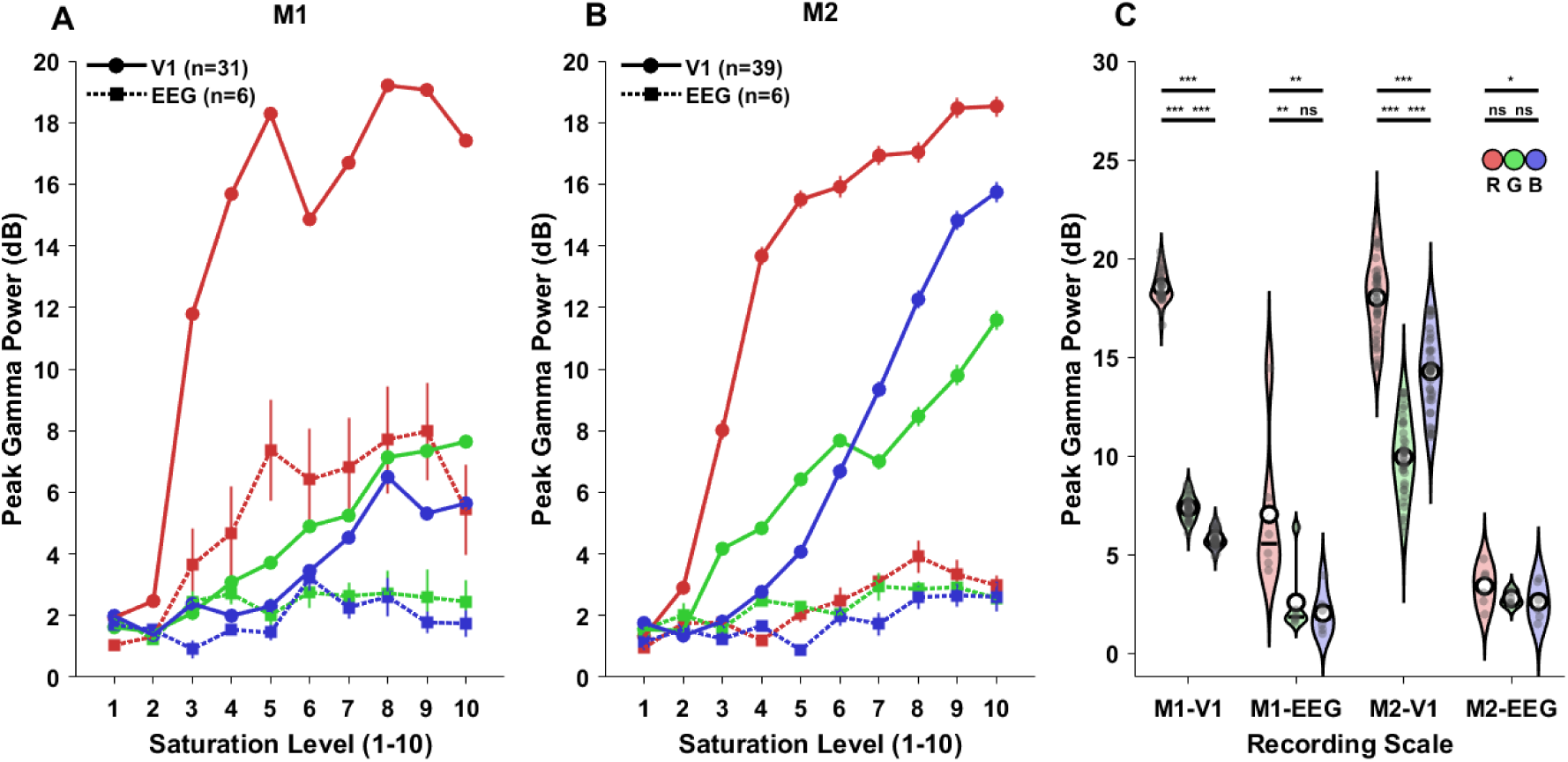
Red hues elicit strongest saturation-dependent gamma in V1 and scalp recordings. Peak gamma power as a function of saturation level (1-10) for the three monitor primaries (red, green, blue) in M1 (A) and M2 (B), with circles showing V1 recordings and squares showing simultaneous scalp EEG measurements. Each data point represents mean ± SEM across recording sites. (C) Population comparison across recording modalities and brain areas shows distributions of peak gamma power for red (R), green (G), and blue (B) primaries at high saturation (levels 8-10). Open circles indicate the mean while horizontal lines indicate median values; circles show individual recording sites. Asterisks denote statistical significance: *** p < 0.001, ** p < 0.01, * p < 0.05; ns, not significant, obtained using paired t-tests to test whether gamma peak power differed between colors within each region.

### Preference for reddish hues is diminished in the DKL space and further reduced in EEG versus LFP

Next, we used DKL color space to probe the underlying cone-opponent mechanisms. We presented 36 hues sampling the DKL isoluminant plane at 10° intervals. Time-frequency analysis showed weaker chromatic selectivity in this space, which further varied across monkeys (Figure 4). Presenting desaturated colors sampled from the DKL space weakened red-selective responses in V1. Monkey 1 (n = 32 sites) generated strong narrowband gamma for long-wavelength hues (0°–60° and 300°–350°), with weaker responses to mid-spectrum greenish hues (120°–180°). Monkey 2 (n = 39 sites), by contrast, showed balanced sensitivity, with gamma responses of comparable magnitude for both reddish and cyan-green hues (180°–240°). Interestingly, simultaneously recorded scalp EEG recordings reflected a further reduction in selectivity compared to V1. This was most evident in Monkey 1, where relatively robust gamma was observed for greenish hues in EEG (n = 4 sites). In Monkey 2 (n = 3 sites), EEG showed similar balanced chromatic sensitivity as seen in V1.

**Figure 4:**
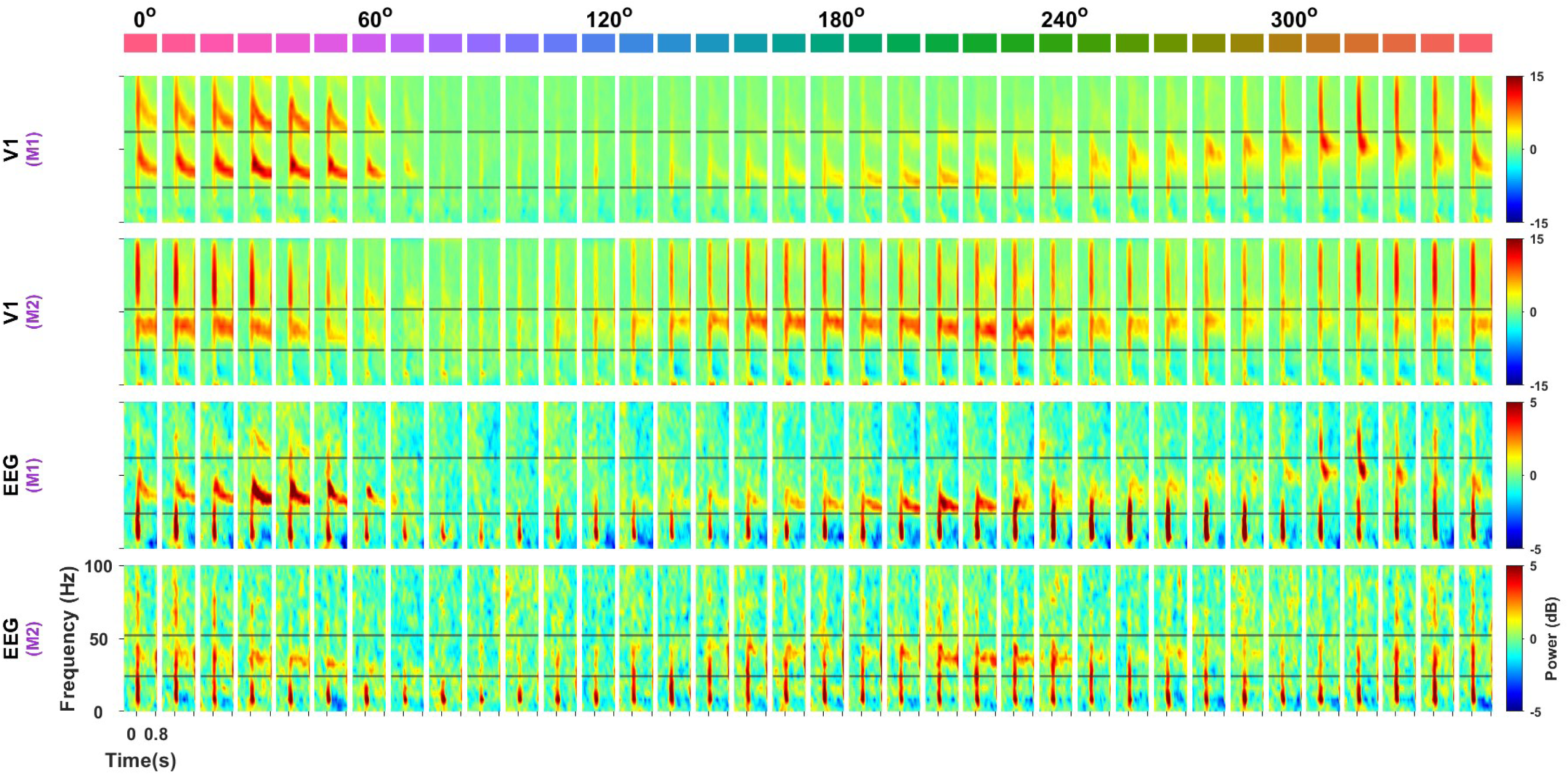
Gamma oscillations induced by colors in DKL space across brain regions and neural scales. Time–frequency spectra showing the decibel change in power from the baseline period (−0.5 s to 0 s, where 0 indicates stimulus onset) for 36 full-field colors sampled around the DKL hue ellipse at equally spaced azimuth angles (0° to 350° in 10° steps; indicated by color swatches at top, where 0°=reddish, 120°=greenish, 240°=bluish). Rows show population averages from V1 LFP (rows 1–2; n = 32 and 39 sites for Monkey 1 (M1) and Monkey 2 (M2), respectively) and scalp EEG (rows 3–4; electrode configuration as described in Methods) recordings from M1 (n=4) and M2 (n=3). Each small panel corresponds to one hue angle. Time axis spans -0.5–0.8 s; frequency axis, 0–100 Hz. Black horizontal lines indicate the gamma band limits used for quantitative analysis.

### L–M cone contrast drives asymmetric gamma modulation in V1

To quantify the chromatic selectivity across scales and probe the specific chromatic mechanisms driving gamma oscillations, we systematically varied color along the L–M (red–green) cardinal axis while keeping the luminance constant (Figure 5). This approach allowed us to isolate the contribution of the red-green opponent pathway. We presented full-field stimuli that ranged from strong negative L–M contrasts (greenish hues, where M-cone activation exceeded L-cone activation) through neutral gray (zero contrast) to strong positive L–M contrasts (reddish hues, where L-cone activation dominated).

**Figure 5:**
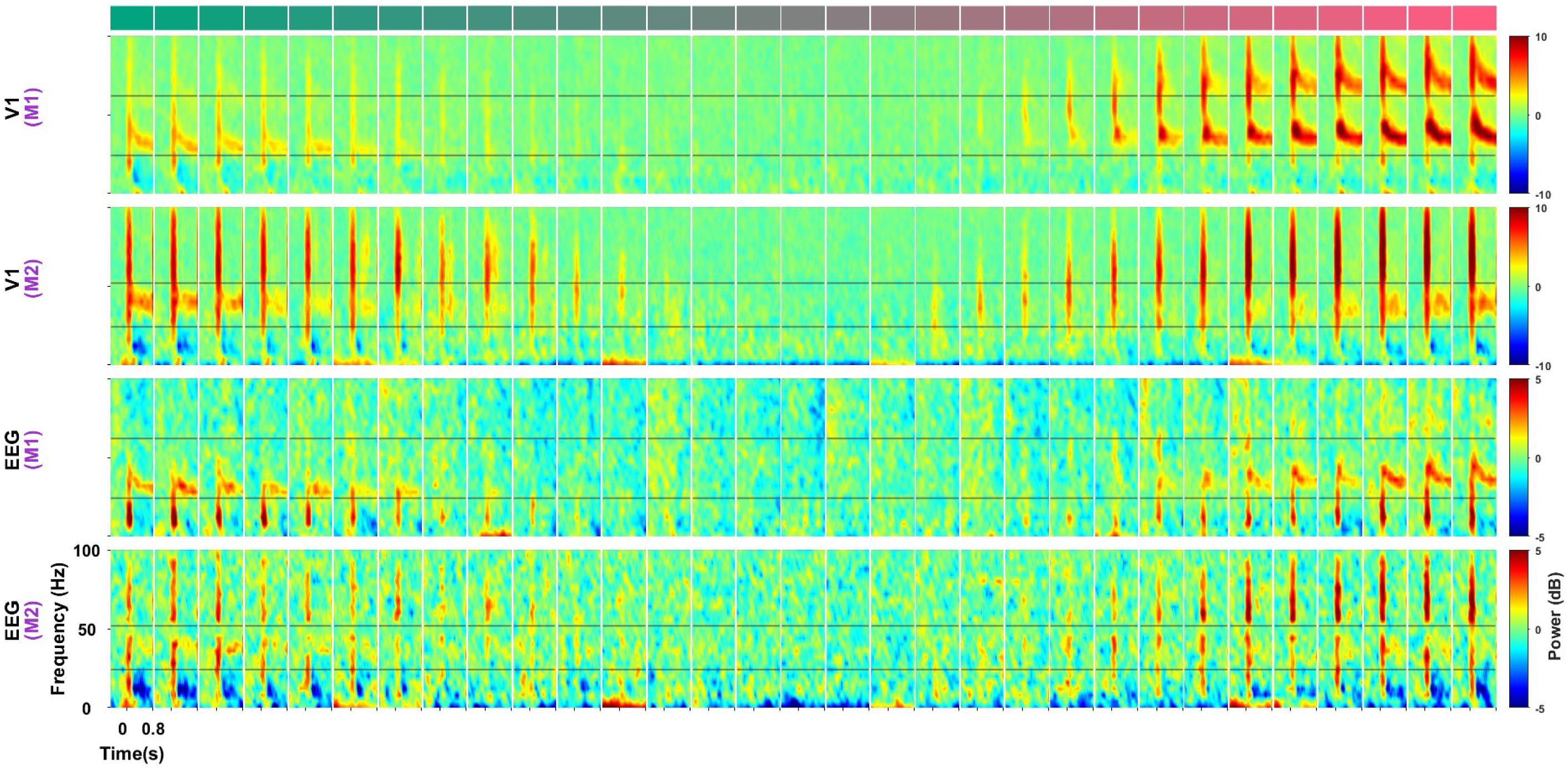
Gamma oscillations scale with L-M cone contrast. Time–frequency spectra showing the decibel change in power from the baseline period (−0.5 s to 0 s, where 0 indicates stimulus onset) for full-field stimuli varying systematically along the L-M cardinal axis at multiple cone contrast levels (indicated by color swatches at top, ranging from negative L-M contrast (greenish, left) through zero contrast (neutral gray, middle) to positive L-M contrast (reddish, right)). All stimuli were isoluminant at 60.27 cd/m². Rows show population averages from V1 local field potential (LFP; rows 1–2) and scalp EEG (rows 3–4) recordings from M1 and M2, respectively. Each small panel corresponds to one L-M contrast level. Time axis spans -0.5–0.8 s; frequency axis, 0–100 Hz. Black horizontal lines indicate the gamma band limits used for quantitative analysis.

In Monkey 1 (n = 32 sites), gamma power increased with L–M contrast in the reddish direction, with stronger responses than equally strong contrasts in the greenish direction. In Monkey 2 (n = 38 sites), V1 gamma responded to L–M contrast in both directions—positive (reddish) and negative (greenish) contrasts of similar magnitude evoked comparable gamma power, consistent with the balanced sensitivity to red and green hues observed in DKL space for this animal. However, the strong V1 red preference observed in Monkey 1 did not persist in simultaneous EEG recordings (M1: n=4, M2: n=3), where neither monkey showed any systematic bias favoring one end of the L–M axis.

We quantified these chromatic asymmetries by measuring peak gamma power in a 0.25–0.75 s window after stimulus onset and computing an “end bias” metric—the difference in gamma power between the strongest positive and negative L–M contrast conditions (Figure 6A–C). This metric captures the degree of asymmetry along the red-green chromatic axis. For Monkey 1, the end bias was significantly different between V1 and EEG (two-tailed independent sample t-test; t (34) = 7.16, p=2.79×10^-8^), confirming the strong asymmetric preference for reddish over greenish stimuli observed in the time-frequency spectra in V1 which was not visible in EEG. In monkey 2, the end bias did not differ between V1 and EEG (two-tailed t-test; t (39) = 0.63, p=0.53), consistent with this monkey’s balanced responses across the L–M axis.

**Figure 6:**
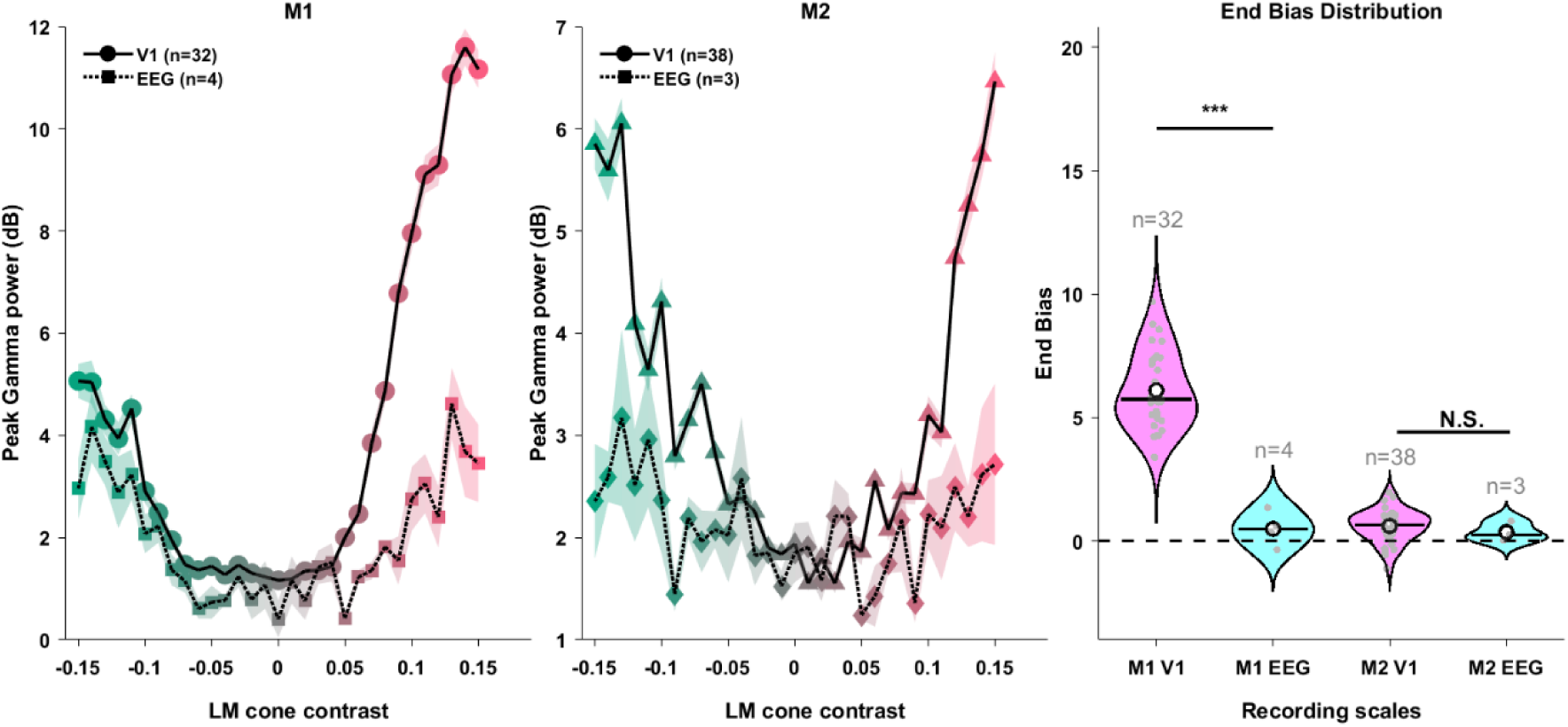
LM cone contrast predicts gamma power and reveals a reddish end bias. Peak gamma power (0.25–0.75 s; dB change from baseline) as a function of L–M cone contrast for M1 (A) and M2 (B). Solid lines, V1 LFP (M1: n=32; M2: n=38); dotted lines, occipital EEG (M1: n=4; M2: n=3). Shaded bands indicate ± SEM. Gamma power shows a pronounced asymmetry in V1 of M1: responses grow steeply for positive L–M contrast (L>M; reddish direction) and remain comparatively weak for negative L–M contrast (M>L; greenish direction). C) “End bias”: the difference in peak gamma power between the positive and negative L–M extremes at matched contrast (∼±0.15). Larger values indicate stronger reddish bias. End bias is significantly greater than zero in M1 V1 (*** p<0.001, two-tailed t-test comparing V1 vs. EEG), but is not different from zero in M1 EEG, M2 V1, or M2 EEG (N.S.). White dot shows the mean while black bar indicates the median; sample sizes are indicated above each distribution.

### Dense color space sampling reveals systematic gamma topography

Finally, by densely sampling stimuli throughout a broad range of the CIE chromaticity diagram, we found that the bias toward long-wavelength colors persisted across continuous variations in hue and saturation (Figure 7). We presented a wide array of colors covering the accessible gamut on our display (with fine increments in saturation and hue) and plotted gamma power responses across this color space for two monkeys. Black curves overlaid on the CIE diagrams in Figure 7 indicate the boundaries of the DKL color space sampled earlier (note that DKL is a relative color space where the cone contrast relative to the pre-stimulus gray background is used to generate the coordinates. Therefore, the DKL colors used in experiment 2 relative to the background gray of ∼60 cd/m^2^ and the isoluminant colors with luminance of ∼8 cd/m2 on the ellipse shown in Figure 7 relative to the background gray of ∼8 cd/m2 have the same coordinates in the DKL space). The resulting gamma power maps showed a strong gradient, with the largest gamma responses confined to the long-wavelength (reddish) region of color space in both V1 and scalp EEG recordings. In addition, we found that colors along the cardinal blue-yellow (S-(L+M)) axis produced negligible gamma power, represented as an elongated blue patch along the S-(L+M) axis. Scalp EEG measurements showed a qualitatively similar color preference topography, albeit with a reduced dynamic range of gamma power changes (approximately 0–8 dB across the color map, compared to 0–20 dB in V1). This shows that the induced gamma is mainly generated by the L-M cone contrast and does not depend on the S cone activation (this was also verified by running an additional experiment where we presented colors across S-(L+M) axis sampled from DKL space, not shown).

**Figure 7:**
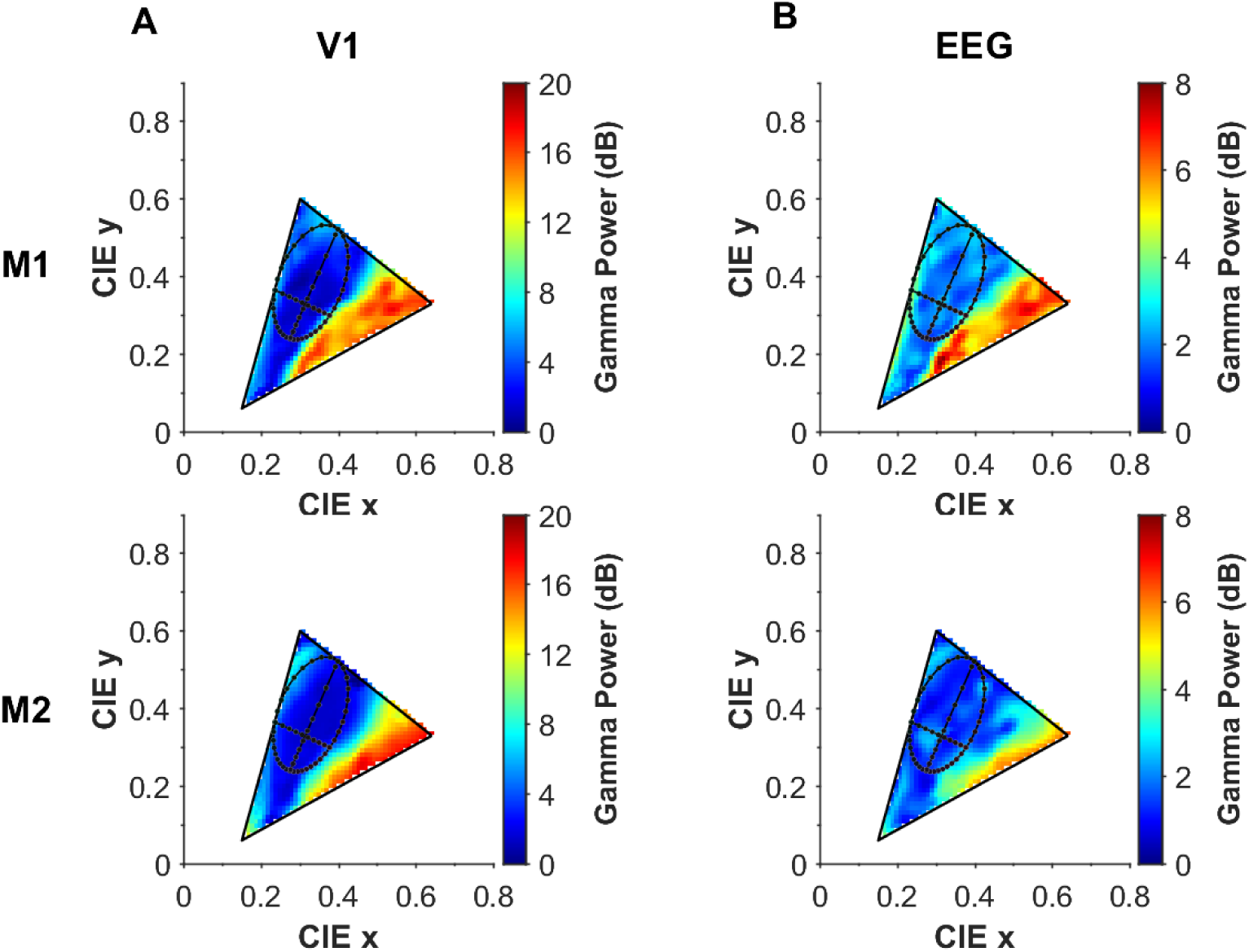
Red bias in gamma oscillations persists across the full CIE color gamut. Gamma power distribution across the CIE chromaticity space for dense radial-angular sampling of the accessible CIE color space at low background luminance (7.86 cd/m²). The black ellipse indicates the boundary of the sampled DKL color space, while the black lines indicate the cardinal axes (L-M and S-(L+M)) overlaid on the CIE chromaticity diagram. (A) Gamma power (25-80 Hz) measured from V1 for monkeys M1 (n=31) (top) and M2 (n=39) (bottom). (B) Gamma power measured from scalp EEG for the same two monkeys (M1: n=4, M2: n=3). Color indicates the change in gamma power (dB) from baseline (-0.5 to 0 s) to stimulus period (0.25-0.75 s). Note the different color scales for V1 (0-20 dB) and EEG (0-8 dB).

In addition to modulating gamma power, chromatic stimuli systematically shifted gamma peak frequency across the CIE color space. Gamma peak frequency was higher towards the long-wavelength (reddish) region (Supplementary Fig. 2). In V1, saturated reddish stimuli elicited peak frequencies reaching ∼60 Hz in M1 and ∼50 Hz in M2, compared to ∼30 Hz for desaturated colors. Scalp recording showed similar trends for both the monkeys, albeit the peak frequency estimation was noisy as gamma peak was not salient for many colors (data not shown).

V4 recordings in Monkey 1 (n = 34 sites) showed weak gamma modulation compared to V1 across all the three experiments (Supplementary Figure 3). In the radial saturation experiment, V4 produced only weak narrowband oscillations at the highest red saturation levels, with negligible responses for green or blue primaries (also note the difference in scale compared to Figure 2). Along the DKL hue ellipse, gamma modulation was virtually absent regardless of hue angle. Similarly, systematic variation along the L–M axis failed to elicit the robust contrast-dependent gamma observed in V1—even the most saturated reddish stimuli produced only weak responses. This near-absence of chromatic gamma in V4 is consistent with previous findings in both macaque extrastriate cortex and human visual areas beyond V1(16, 29, 32).

## Discussion

We report three major findings. First, saturated reddish hues consistently evoked the strongest V1 gamma, but this response scaled monotonically with L–M cone contrast and was virtually absent along the S-cone axis—demonstrating that L–M drive, rather than hue category, is the primary determinant of color-induced gamma. Second, this robust V1 signal did not propagate uniformly across neural scales: V4 showed minimal color-induced gamma, and chromatic selectivity was attenuated in scalp EEG. Third, gamma power reduced with desaturation, such that selectivity for reddish hues diminished when colors were constrained to low-saturation DKL ellipses. Together, these results reconcile invasive and non-invasive findings and demonstrate that “reddish” colors are potent gamma drivers because they maximize L–M cone contrast—and that this oscillatory signal is primarily observed in early visual cortex.

### Reconciling the “red bias” with cone-contrast accounts

Our data show that the “red bias” and “cone-contrast dependence” describe the same underlying mechanism. Early invasive work demonstrated that large, uniform reddish stimuli evoke unusually strong narrowband gamma in macaque V1 (16, 28, 29, 33). Human ECoG and MEG studies confirmed that uniform color fields induce strong gamma, with red–orange stimuli typically producing the largest responses (34–36). However, the precise mechanism has remained contested because most studies used colors defined in CIELAB/CIELUV or monitor-based spaces that are matched for lightness but do not guarantee physiological isoluminance or equated cone contrast. Hue, saturation, and L–M cone contrast are thus partially confounded: “red” stimuli occupy regions of color space with relatively high L–M contrast, whereas “green” and “blue” stimuli have weaker L–M drive. This issue was addressed by Stauch et al. (2022) using stimulus generated in DKL space in human MEG: when L–M cone contrast was equated, the red-over-green preference was not observed (32). Our simultaneous V1–V4–EEG recordings extend these observations. By sampling colors from both from the CIE and DKL color space, we show that L–M contrast, rather than hue category, primarily governs V1 gamma magnitude. Highly saturated reddish stimuli have high L–M contrasts which drive V1 gamma strongly. Constraining stimuli to a fixed-radius DKL ellipse reduced overall cone contrast, weakening gamma power and reducing the reddish hue bias—consistent with prior findings (32).

### Mechanisms: E–I circuitry and cone-opponent pathways

Narrowband gamma emerges from recurrent excitatory–inhibitory (E–I) networks driven into a PING-like regime by sufficient excitatory input (1–3). Recent work shows that gamma evoked by hues and achromatic gratings share a characteristic non-sinusoidal waveform, reproducible only when a Wilson–Cowan network is inhibition-stabilized with superlinear inhibition (37). This supports a shared PING-like mechanism for both luminance- and color-induced gamma.

Our results extend this picture by showing that cone-opponent architecture tightly constrains which chromatic directions engage this circuitry. Cone-opponent signals originates in the retina and LGN and then undergo further spatial processing in V1 through single- and double-opponent neurons clustered in cytochrome-oxidase blobs (38–40). Uniform L–M surfaces provide spatially coherent excitation and surround inhibition, creating ideal conditions for robust interneuron recruitment.

The striking L–M versus S-cone difference reflects fundamental pathway asymmetries. The L–M pathway via the parvocellular system provides dense, temporally precise input to layer 4Cβ. The S-cone pathway via the koniocellular system has distinct cortical terminations (layers 2/3 and 4A), slower dynamics, and sparser connectivity (39, 41). It is therefore plausible that koniocellular drive is too weak and temporally dispersed to robustly entrain fast-spiking interneurons, consistent with very weak S-cone–driven gamma observed in both our results and previous human MEG study (32). Within the L–M axis, the “end bias” favoring reddish stimuli in both the monkeys may reflect the ∼2:1 L:M cone ratio in macaque fovea (42) or differential adaptation rates between cone pathways (28). The individual variability—Monkey 2 showed more balanced L–M sensitivity—suggests asymmetry may vary with cortical location or cone-opponent wiring. Furthermore the upward shift in peak gamma frequency toward reddish hues (Supplementary Figure 2), parallels the established modulation of oscillatory frequency by luminance contrast (22, 43). In Wilson-Cowan models, such frequency shifts reflect stronger excitatory drive accelerating E–I loop dynamics (37, 44), suggesting that saturated L–M stimuli recruit gamma-generating interneuron networks more effectively than desaturated colors.

### Why V4 and scalp EEG show weaker color-induced gamma?

The near absence of narrowband gamma in V4, despite robust V1 responses, aligns with Rols et al. (2001) and MEG source analyses localizing color-driven gamma to early visual cortex (29, 32). This likely reflects a transformation in color representation along the ventral pathway. Early visual cortex (V1) is strongly driven by cone-opponent “cardinal’ axes inherited from the LGN, whereas V4 “globs” contain neurons tuned to perceptual hues arranged in a nearly uniform non-cardinal space (40, 45, 46). Large uniform color fields recruit many V4 modules with different preferred hues; when pooled, these diverse sources may partially cancel, yielding weak narrowband gamma even when firing rates remain robust.

Although V1 gamma showed red bias, scalp EEG responses were symmetric or showed weaker red preferences. This could be because EEG integrates signals from multiple visual areas (V1,V4) with heterogeneous chromatic tuning, which could weaken the asymmetries evident in local V1 recordings (28, 32).

### Implications for predictive coding

Gamma has been proposed to carry feedforward prediction errors (11, 12, 47). In our experiments, we used highly predictable full-field surfaces, yet saturated L–M stimuli produced enormous gamma while S-cone stimuli—despite being visible—produced almost none. If gamma directly encoded prediction error, comparable responses would be expected regardless of chromatic content once contrast is controlled.

A more parsimonious interpretation is that gamma reflects the degree to which a cortical population engages in recurrent E–I computation. The parvocellular pathway recruits gamma-generating circuitry effectively; the koniocellular pathway does not. Future work should examine whether prediction errors along sparsely represented channels rely on different neural codes, and how predictive coding models might accommodate such pathway-dependent constraints.

### Clinical relevance

Red and red–blue flickers trigger photoparoxysmal responses in photosensitive epilepsy, whereas blue–green flickers are less epileptogenic (48, 49). This pattern mirrors gamma’s stimulus tuning. Although correlational, it raises the possibility that strong L–M-driven gamma may predispose susceptible individuals to pathological hypersynchrony (35, 50). Also, visually induced gamma has emerged as a biomarker for aging and neurodegeneration—gamma power declines with age (5, 8), is reduced in mild cognitive impairment and Alzheimer’s disease (4), and correlates with cognitive status (51) These studies used achromatic stimuli; given that chromatic gamma could reflect E–I circuit dynamics known to be disrupted in aging and disease, future studies should explore whether chromatic gamma signatures offer additional sensitivity or specificity as biomarkers for aging and neurodegenerative disease.

## Author contributions

Conceptualization: AB, SR. Methodology: AB. Supervision: SR. Writing: AB, SR

## Acknowledgments

We thank Prof. Bevil Conway, Prof. David Brainard, Prof. John Maunsell, Prof. Geoffrey Ghose for their guidance on DKL color space characterization. Dr. Vinay Shirhatti for initial project guidance; Dr. Divya Gulati (EEG electrode layout) and Dr. Surya Prakash for helpful discussions and assistance with recordings; and Sveekruth Pai, Gouroju Shashank (display calibration, MonkeyLogic Rig setup, and recordings), and Niloy Maity (recordings) for technical assistance.

## Materials and Methods

### Animal Preparation and training

All procedures followed protocols approved by the Institutional Animal Ethics Committee (IAEC) at the Indian Institute of Science and complied with Committee for the Purpose of Control and Supervision of Experiments on Animals (CPCSEA) guidelines. We worked with two adult macaque monkeys (*Macaca radiata*; M1: female, 15 years, ∼5.8 kg; M2: male, 17 years, ∼7.2 kg). Under general anesthesia, we first implanted a titanium headpost on each animal’s skull. After recovery, we trained the monkeys to perform a passive visual fixation task. Once training was complete, we performed a second surgery under general anesthesia to implant dual 48-channel Utah microelectrode arrays (1-mm electrode length; 400-µm spacing; Blackrock Microsystems) in the right hemisphere, targeting V1 (10–15 mm anterior to the occipital ridge, 10–15 mm lateral to the midline) and V4 (35 mm anterior, 35 mm lateral). Minor placement variations occurred across animals. Reliable V4 responses were obtained only from Monkey 1; in the second monkey, the V4 arrays did not yield reliable single units or visually evoked responses, so V4 analyses were restricted to Monkey 1.

Recorded neurons from V1 had receptive fields in the lower-left visual quadrant at eccentricities of ∼2.6°–3.3° in M1 and ∼2.8°–3.5° in M2. V4 receptive fields in M1 were at eccentricities of ∼1.0°–2.2°. Given the array size relative to cytochrome-oxidase blob dimensions, our V1 recordings likely sampled both blob and interblob compartments. The units displayed orientation tuning (data not shown), consistent with substantial sampling from interblob regions where orientation selectivity is pronounced.

We also recorded 18-channel EEG using Ag–AgCl cup electrodes (Grass Technologies) positioned on the scalp (Refer to Supplementary Figure 1 for the electrode montage). Before each session, we trimmed the hair and prepared the scalp with a mild abrasive gel (NuPrep Skin Prep Gel, Weaver and Company) to reduce impedance, then applied Ten20 Conductive Paste (Weaver and Company) to secure the electrodes. As sessions lasted 3–4 hours, electrodes were additionally secured with Medi-tape strips to prevent movement. The ground electrode was placed frontally, with hardware grounding via the Digital Hub connected to a dedicated earth line.

### Experimental setup and behavior

During behavioral sessions, each monkey sat in a primate chair with its head stabilized using the implanted headpost. Visual stimuli were presented on a calibrated LCD monitor (BenQ XL2411; 1280 × 720 resolution; 100 Hz refresh rate) positioned approximately 50 cm from the eyes. The experimental setup was housed within a Faraday enclosure with dedicated grounding isolated from the mains supply.

The monkeys performed a passive fixation task. Each trial began with fixation onset on a small central point (0.10° radius), followed by a 1000-ms blank gray screen. We then presented 2–3 visual stimuli, each shown for 800 ms with 700-ms interstimulus intervals. Animals received a juice reward for maintaining fixation within 1.5° around the fixation point throughout the trial.

### Monitor calibration

All stimuli were presented on a BENQ XL2411 LCD monitor. The display was gamma-corrected (γ = 1.0) for each primary using an i1Display Pro colorimeter, which was used to generate a linearized ICC profile. The ICC profile reported the following CIE xy chromaticities for the primaries and white points: Red (0.644, 0.329), Green (0.325, 0.606), Blue (0.160, 0.065), and white (0.346, 0.358). Because colorimeters estimate chromaticity using model-based approximations rather than direct spectral measurements, we re-measured the primaries and the white point with a PR-655 spectroradiometer after loading the ICC profile onto the monitor. The spectroradiometer values were: Red (0.641, 0.333), Green (0.309, 0.616), Blue (0.157, 0.071), and white (0.323, 0.348). All subsequent calculations used these spectroradiometer values. Detailed explanation of the color space calculations is provided in Appendix 1 and 2.

### Data recording

We recorded neural activity from 96 channels using a 128-channel Cerebus Neural Signal Processor (Blackrock Microsystems). Our dual 48-channel Utah microelectrode arrays were connected to the first two inputs of the Digital Hub, occupying channels 1–96 (banks 1–3). Local field potentials were obtained by applying online filtering to the raw signals using a first-order analog Butterworth high-pass filter at 0.3 Hz and a fourth-order digital Butterworth low-pass filter at 500 Hz. The filtered signals were sampled at 2 kHz with 16-bit resolution. For spike activity, we applied a separate online filtering cascade (250-Hz high-pass, fourth-order digital Butterworth; 7.5-kHz low-pass, third-order analog Butterworth) and detected threshold crossings at approximately five standard deviations above baseline noise. All subsequent analyses were performed on these signals without additional offline filtering. We also recorded concurrent 18-channel scalp EEG (Supplementary Figure 1) through the fourth bank (channels 97–128) using a breakout box (Omnetics, Blackrock Microsystems). This configuration ensured perfect temporal synchronization between intracortical and scalp recordings within the same data stream.

Eye position (horizontal and vertical coordinates) and pupil diameter were monitored throughout each session using an ETL-200 Primate Eye Tracking System (ISCAN Incorporated). Task control, stimulus generation, and pseudorandomized stimulus presentation were implemented using MonkeyLogic running on a Windows workstation.

### Electrode selection

We selected electrodes based on signal quality and receptive field stability. To map receptive fields, we presented small sinusoidal gratings across a rectangular grid spanning the array’s aggregate receptive field and analyzed evoked response potentials (ERPs) to identify sites with reliable stimulus-driven responses. Repeating these procedures across multiple sessions allowed us to identify electrodes with consistent stimulus responses and stable receptive field properties. This procedure resulted in 32 electrodes for M1 V1, 34 electrodes for M1 V4 and 39 electrodes for M2 V1. Electrodes with impedances <100 kΩ or above upper thresholds (M1: >2500 kΩ; M2: >3000 kΩ) were excluded for each recording session, resulting in 31-32 electrodes for M1 and 38-39 electrodes for M2 for different experiments.

For EEG analysis, we excluded channels with high electrode–skin impedance (> 25 kΩ) and considered only electrodes that showed a strong stimulus-locked ERP. Applying these criteria on each recording session yielded 3–6 EEG electrodes per monkey, which were predominantly located over the occipital region.

### Data Analysis

We analyzed local field potential (LFP) data using multitaper spectral methods implemented in the Chronux toolbox (http://chronux.org), a MATLAB-based package optimized for neurophysiological time-series analysis. Time-frequency spectrograms were computed using a single taper (time-bandwidth product = 1, number of tapers = 1) with a 250 ms sliding window, yielding a frequency resolution of 4 Hz—sufficient to resolve gamma-band dynamics while maintaining temporal precision. We computed power spectral density (PSD) during two epochs: “baseline” period (−0.5 to 0 s before stimulus onset) and “stimulus” period (0.25 to 0.75 s after onset). We excluded line noise at 50 Hz and its harmonics from all analyses.

### Gamma power

For analyses of gamma power, we defined the gamma band separately for the CIE and DKL stimulus sets. For CIE stimuli, we used a broader band (25–80 Hz) because peak gamma frequency varied substantially across conditions and sessions, and this range captured the response consistently in both monkeys. For DKL stimuli, peak gamma frequency was more narrowly distributed; we therefore used narrower, monkey-specific bands (M1: 24–62 Hz; M2: 24–52 Hz) to match the observed peak range.

To quantify stimulus-driven changes, we expressed gamma power relative to baseline in decibels:

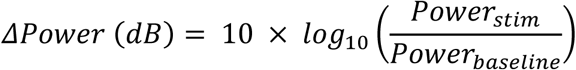

This logarithmic transformation captures the multiplicative nature of power changes and normalizes the distribution of gamma responses across stimulus conditions. We employed a common baseline approach, pooling trials across all stimulus conditions to compute a stable baseline reference. This method avoids potential biases from condition-specific baseline fluctuations and ensures consistent comparison across different chromatic stimuli.

### Peak Gamma frequency and power

For analyses quantifying maximal responses, we defined peak gamma frequency as the frequency corresponding to the maximum relative power change within the gamma frequency band and power (*P*_peak_) as value of the power change at this peak frequency:

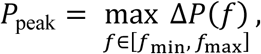

where Δ*P*(*f*)denotes the stimulus-induced change in power spectral density at frequency *f* (in dB relative to baseline), and [*f*_min_, *f*_max_] specifies the gamma-band frequency range. For estimating the peak frequency, we only considered frequencies within the gamma frequency band for which gamma power was more than 5 dB.

### Population-Level Analysis

Gamma power was calculated independently for each electrode. For population analyses, we averaged gamma power across electrodes within each recording area (V1, V4, EEG) and calculated population statistics as mean ± SEM, where SEM is the standard error of the mean (σ/√n).

## Supplementary Material

**Supplementary Figure 1:**
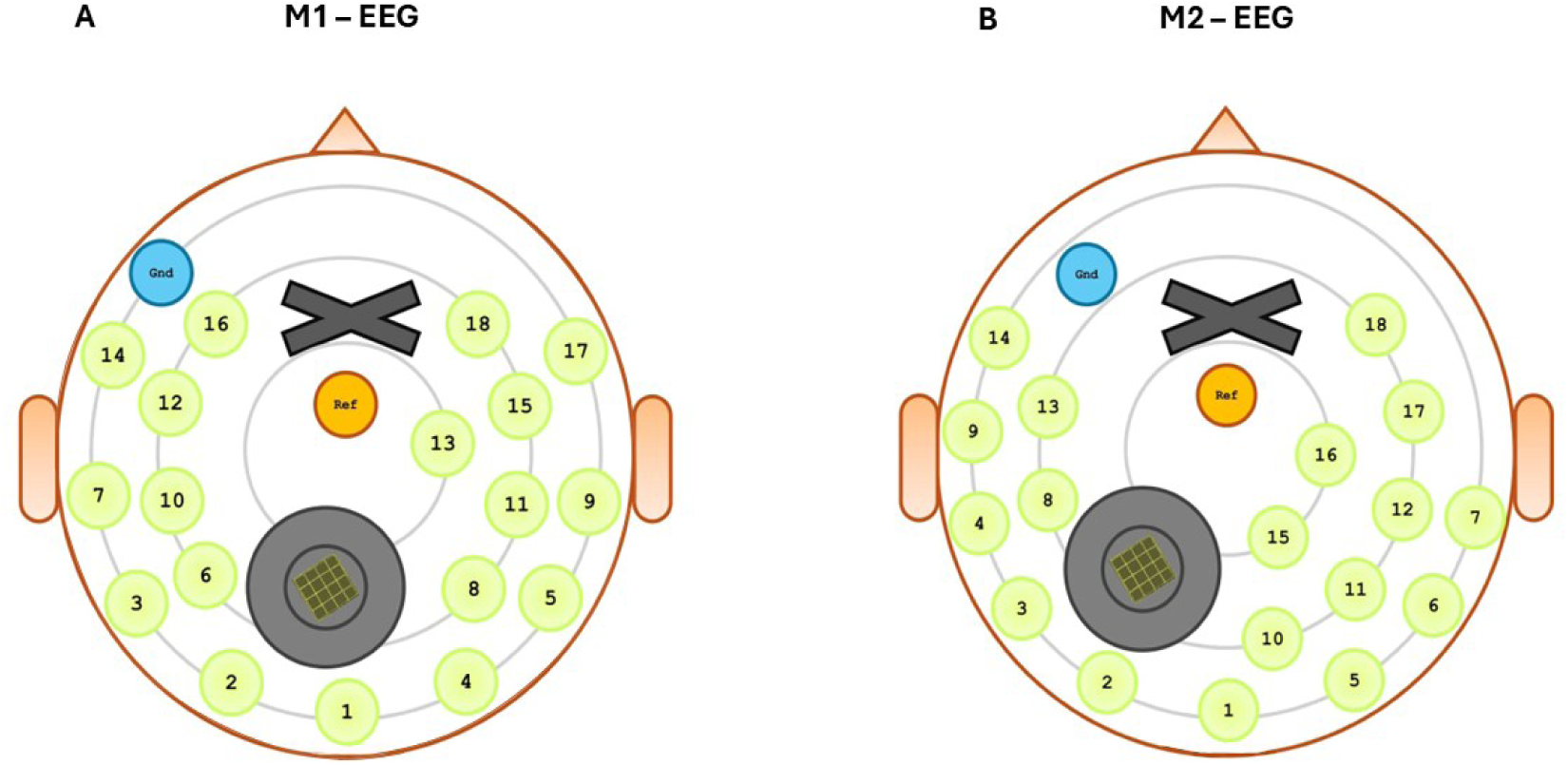
EEG electrode montage for M1 and M2. Schematic of 18-channel EEG electrode positions for M1 (A) and M2 (B), shown from a dorsal view. Green circles indicate recording electrodes (numbered 1–18). The ground electrode (Gnd; blue) was placed at a frontal location. The reference electrode (Ref; orange) was placed at a central location near the midline. The gray region with crosshatch pattern indicates the location of the connector. The X marks the headpost location. Exact electrode positions varied slightly between animals depending on headpost and connector placement. Analyses focused on occipital electrodes showing robust visually evoked responses.

**Supplementary Figure 2:**
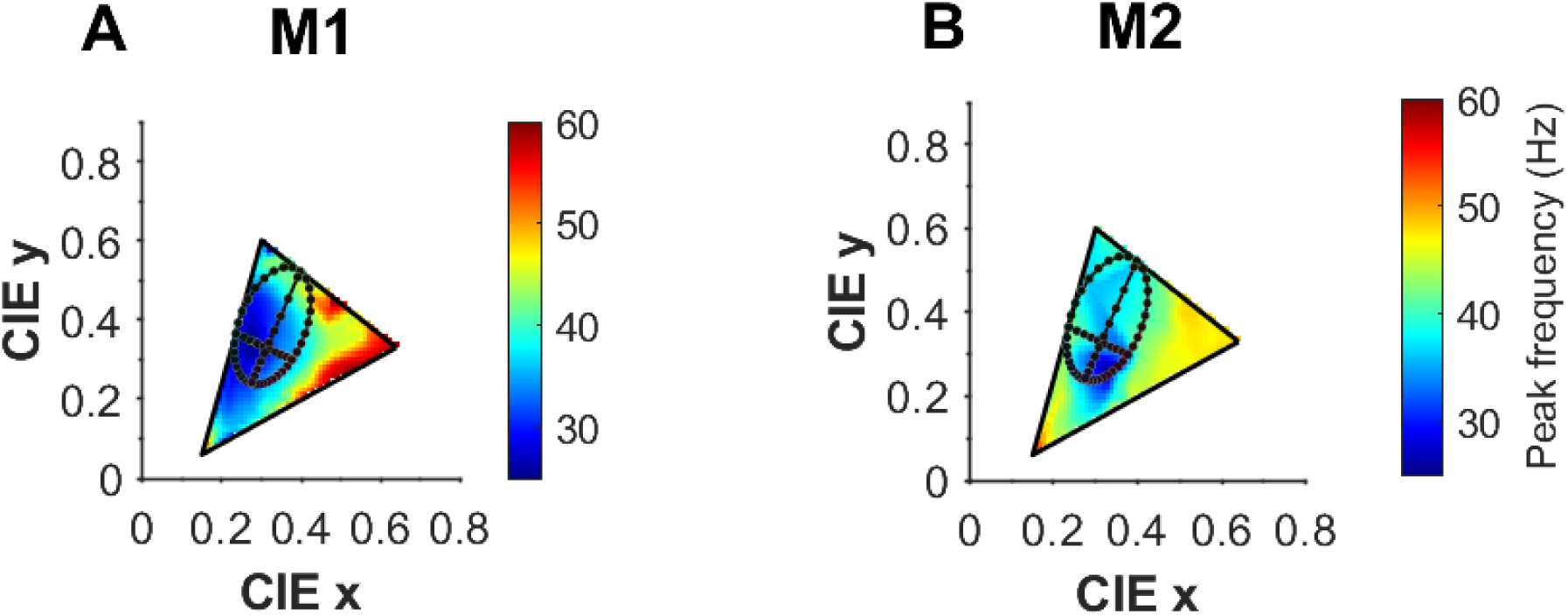
Peak gamma frequency increases for long-wavelength hues across the CIE color gamut. Peak gamma frequency distribution across the CIE chromaticity space for dense radial-angular sampling of the accessible CIE color space at low background luminance (7.86 cd/m²). The black ellipse indicates the boundary of the sampled DKL color space, while the black lines indicate the cardinal axes (L-M and S-(L+M)) overlaid on the CIE chromaticity diagram. a) Peak gamma frequency measured from V1 for M1 (A) and M2 (B). Color indicates the frequency (Hz) at which gamma power reached its maximum during the stimulus period (0.25–0.75 s).

**Supplementary Figure 3:**
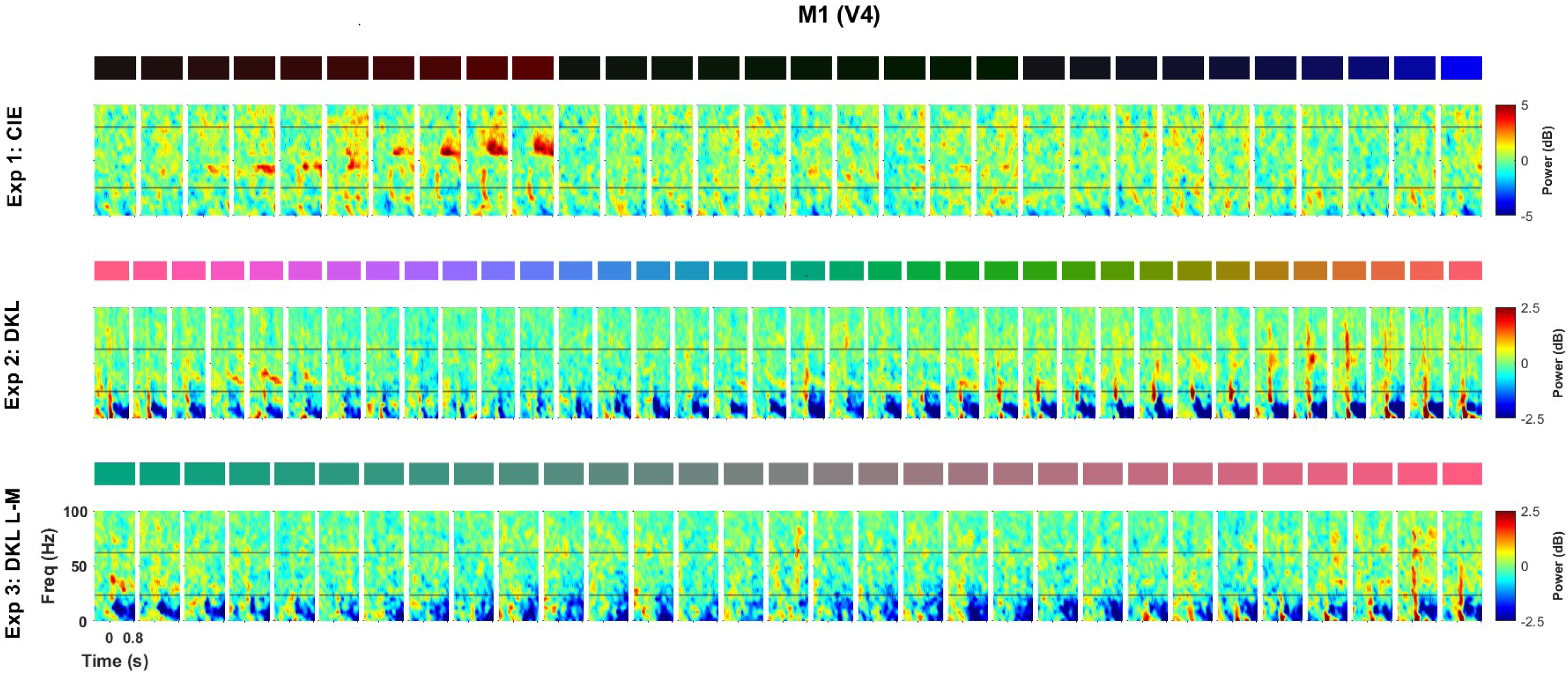
V4 shows minimal color-induced gamma across all three different experiments. Time–frequency spectra showing the decibel change in power from the baseline period (−0.5 s to 0 s, where 0 indicates stimulus onset) for area V4 in Monkey 1 (n = 34 sites) across three different experiments. **Top row (CIE):** Radial saturation variation toward the three monitor primaries (red, green, blue) at 10 saturation levels each (indicated by color swatches; progressing from dark/desaturated to saturated). **Middle row (DKL Hue):** 36 full-field colors sampled around the DKL hue ellipse at equally spaced azimuth angles (0° to 350° in 10° steps; 0°=reddish, 120°=greenish, 240°=bluish). **Bottom row (DKL L-M):** Stimuli varying systematically along the L-M cardinal axis from negative L-M contrast (greenish, left) through zero contrast (gray, center) to positive L-M contrast (reddish, right). Note the compressed color scales (±5 dB for CIE; ±2.5 dB for DKL experiments) compared to V1.

## Appendix 1

### Visual Stimuli

We designed four experiments to systematically characterize how chromatic inputs modulate gamma oscillations. All stimuli appeared as full-field color patches on the calibrated LCD monitor. We controlled stimulus timing and task flow using custom MATLAB scripts using the MonkeyLogic toolbox. Each trial comprised an 800-ms stimulus presentation followed by a 700-ms inter-stimulus interval during which the display returned to the background luminance. Stimuli were presented in pseudorandom order to minimize adaptation effects. We used full-field stimuli because gamma oscillations, unlike single-neuron responses, increase in strength with stimulus size and are most robust for large-field stimuli.

### Experiment 1: Radial saturation variation toward monitor primaries

For each primary direction (red, green, blue), we calculated the theoretical chromaticity coordinates along a line from the white point to that primary, generated 10 equally spaced steps along each line in xyY space with Y fixed at our target value, then transformed these xyY specifications to RGB values for display. We measured the spectra of all the stimuli with the PR-655 spectroradiometer. The measured luminances were around 7.87 cd/m² across all saturation levels for all three primary directions (Fig. 1A, right panel). The near-constant luminance across the full saturation range—from the gray point to the most saturated colors our display could generate—ensured that saturation was effectively isolated as an independent variable.

### Experiment 2: Isoluminant hue ellipse

We sampled 36 colors uniformly around the DKL isoluminant plane at 10° azimuth intervals (0°–350°, where 0° corresponds to the positive L-M axis). Because monitor gamut constraints produce an elliptical boundary in DKL space—with greater attainable cone contrast along the S-(L+M) axis than the L-M axis—we sampled along the largest ellipse displayable at a background luminance of 59.45 cd/m². Note that the maximum allowable |L-M| contrast depends on the point where the L-M axis intersects the line joining the green and blue primaries, while the maximum value along the S-(L+M) axis depends on the point where this cardinal axis intersects the line connecting the green and red primaries, as shown in Figure 1B on the CIE xy plot. The size of this ellipse does not change if the luminance of the colors is set to a lower value. In fact, hues on the ellipse relative to the white point (pre-stimulus background) at the same luminance level (for example, the ellipse shown in Figure 7) have the same coordinates in the DKL space irrespective of the luminance since DKL is a relative color space.

We measured all the stimulus with the PR-655 spectroradiometer and computed their LMS cone activations, MacLeod-Boynton coordinates (see below for details), and DKL values using our monitor’s calibration data. The measured luminance clustered around the background value of 59.45 cd/m² (Fig. 1B, right panel), confirming that chromatic modulation occurred with negligible residual luminance variation across the entire hue ellipse.

### Experiment 3: L-M cone contrast variation

To probe responses along the cardinal L-M opponent axis, we generated isoluminant stimuli varying systematically from negative L-M contrast (greenish) through zero contrast (background) to positive L-M contrast (reddish). We designed these stimuli in DKL space by keeping the S-(L+M) coordinate and luminance constant while varying the L-M coordinate across different levels (Figure 1C). The background luminance was 60.27 cd/m².

### Experiment 4: Dense color space sampling

We extended the isoluminant saturation approach to map gamma responses comprehensively across accessible color space. With Y fixed at 8.5 cd/m², we sampled extensively within the triangle formed by our monitor’s three primaries in the xy chromaticity plane.

We generated a stimulus grid that sampled both radial paths toward the monitor primaries and intermediate directions, providing dense angular–radial coverage of the displayable color space. This approach included colors along the edges of the monitor gamut triangle, interior regions formed by mixtures of all three primaries, and intermediate points—all at constant luminance. To ensure feasibility, we computed theoretical xyY coordinates and retained only those who’s corresponding RGB values fell within the valid range (0–1), thereby restricting the final stimulus set to colors within the physical display gamut.

Luminance for all presented colors was verified using a spectroradiometer and remained close to 7.86 cd/m² (Fig. 1D, right panel). This isoluminant sampling provided a systematic map of the accessible color space, enabling assessment of gamma responses across the full chromatic plane rather than along isolated axes.

### Color Space Framework and Transformations

Our experiments required stimuli defined in physiologically meaningful color spaces while maintaining precise control over luminance. Monitors produce colors by combining their red, green, and blue primaries (RGB), but these device-dependent values do not directly relate to how the visual system encodes color. To relate device output to visual processing, we transformed RGB values through a series of intermediate color spaces. CIE XYZ provides a standardized, device-independent description of color based on human color matching functions. From XYZ, we computed LMS coordinates—the excitation levels of the long-, medium-, and short-wavelength sensitive cone classes—linking our stimuli to photoreceptor responses. Finally, we expressed colors in DKL space (Derrington, Krauskopf, & Lennie, 1984), which recombines cone signals along three cardinal axes: luminance (L+M), red-green chromatic (L−M), and blue-yellow chromatic (S−(L+M)). These axes correspond to the response properties of distinct neural pathways from retina through lateral geniculate nucleus. We describe each transformation below, with the complete mathematical derivation of the DKL normalization provided in Appendix 2.

### RGB to XYZ Transformation

The first transformation converts from the monitor’s device-dependent RGB space to the device-independent CIE XYZ color space. This transformation requires only the chromaticity coordinates (x, y) of the three primaries and the white point. The 3×3 transformation matrix M_RGB→XYZ emerges from the requirement that the three primaries, when combined in appropriate proportions, produce the target white point. We computed this matrix using the RGBToXYZMatrix function in Psychtoolbox.

### XYZ to LMS Transformation

LMS cone excitation values are obtained from spectral power distributions by integrating with the cone fundamentals. However, standard cone fundamentals are normalized to have a maximum value of 1, so the resulting LMS values require rescaling to ensure that L + M equals photometric luminance Y (the “luminance units” convention). This scaling is essential for the physiological interpretation of DKL coordinates.

The transformation depends on the choice of the cone fundamentals and color matching functions; we used the ones described by Stockman and Sharpe (2000). The relationship between the Stockman & Sharpe (2000) cone fundamentals and the CIE XYZ color matching functions is given by the transformation matrix (Stockman, 2019):

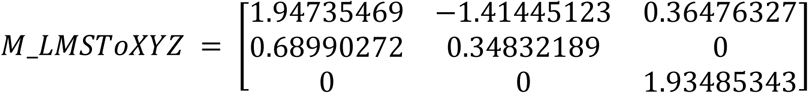

The second row of this matrix provides the luminance weights: Y = 0.6899L + 0.3483M. To convert LMS values to luminance units where L + M = Y, we premultiply the inverse of M_LMS→XYZ by a diagonal scaling matrix with entries [0.6899, 0.3483, 0.0372]. The resulting XYZ-to-LMS transformation matrix in luminance units is:

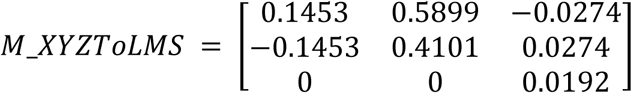

### LMS to DKL Transformation

The DKL color space (Derrington, Krauskopf, & Lennie, 1984) represents color along three physiologically motivated axes corresponding to the response properties of neurons in the lateral geniculate nucleus: an achromatic luminance axis (L+M), a red-green chromatic axis (L−M), and a blue-yellow chromatic axis (S−(L+M)). This space is ideal for isolating specific chromatic mechanisms because modulation along each cardinal axis selectively activates one mechanism while leaving the others unchanged.

The transformation from LMS to DKL coordinates can be expressed as a single matrix operation. Let the LMS coordinates of the background (white point) be [b₁, b₂, b₃]^T^, where b₁ and b₂ are in luminance units such that L = b₁ + b₂. The base transformation matrix B captures each mechanism:

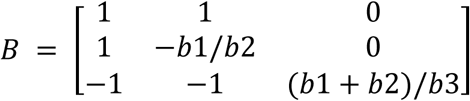

Post-multiplying B by an LMS vector [L, M, S]^T^ yields a vector whose entries are: (1) luminance L+M, (2) the L−M opponent signal scaled by baseline cone excitations, and (3) the S−(L+M) opponent signal similarly scaled.

After multiplying by B, the resultant matrix needs to be appropriately normalized to ensure that each mechanism-isolating stimulus with unit pooled cone contrast produces unit responses in the three DKL mechanisms. The original approach used by Brainard (implemented in both Psychtoolbox’s computeDKL_M function and Westland’s lms2dkl MATLAB toolbox (52) performed these normalizations explicitly for each column. We show in Appendix 2 that this can be achieved by simply pre-multiplying B with a diagonal normalization matrix D:

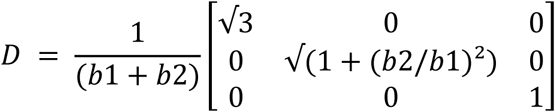

where L = b₁ + b₂. The complete transformation from LMS (in luminance units) to DKL coordinates is thus:

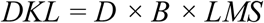

Psychtoolbox’s computeDKL_M function describes our alternate approach in a commented-out section for comparison. Both methods yield identical values in the DKL space.

### MacLeod-Boynton Chromaticity Space

For practical stimulus generation on the isoluminant plane, we used the MacLeod-Boynton (MB) chromaticity diagram (MacLeod & Boynton, 1979), which projects the three-dimensional LMS cone space onto the L+M = 1 plane. The MB coordinates are:

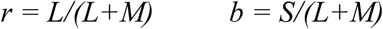

The r coordinate (often called the “constant-blue” or cb axis) represents the L-to-M ratio: moving along this axis changes L and M cone excitations in opposition while keeping S constant. The b coordinate (the “tritanopic confusion” or tc axis) represents S-cone excitation relative to total L+M luminance: moving along this axis changes only S while maintaining a constant L/M ratio.

For DKL stimuli on the isoluminant plane (elevation θ = 0°), the cb and tc axes correspond directly to the L−M and S−(L+M) chromatic directions, respectively. Our procedure for generating isoluminant DKL stimuli was: (1) find the largest ellipse in MB space that fits within the monitor’s displayable gamut; (2) sample points around this ellipse at desired azimuth angles φ; (3) convert each (r, b) coordinate to CIE xy chromaticity; (4) set luminance Y to the desired background value; (5) transform the resulting xyY coordinates to XYZ, then to RGB using the calibrated monitor matrix.

## Appendix 2

### Derivation of the DKL Normalization Matrix

This appendix provides the complete mathematical derivation of the diagonal normalization matrix D used in the LMS-to-DKL transformation. The derivation follows Brainard (1996) and demonstrates that D can be computed directly from the background LMS coordinates without iterative procedures.

### Problem Statement

Given the base transformation matrix B (defined in Methods), we seek a diagonal matrix D such that the complete transformation DKL = D × B × LMS satisfies the following normalization condition: each mechanism-isolating stimulus with unit pooled cone contrast produces unit responses in the corresponding DKL mechanism.

*Below, we show that the diagonal normalization matrix D can be computed directly as:*

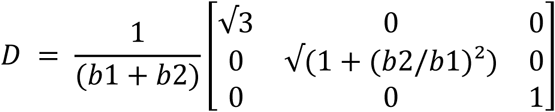

*where L = b₁ + b₂ is the background luminance, and [b₁, b₂, b₃]^*T*^ are the LMS coordinates of the background in luminance units*.

### Proof

The derivation proceeds in five steps.

#### Step 1: Invert the base matrix B

The inverse of B provides the three mechanism-isolating stimuli as its columns. Since B has a simple structure (only two entries differ from 0 or 1), we can compute the inverse directly by the adjoint-determinant method. The result is:

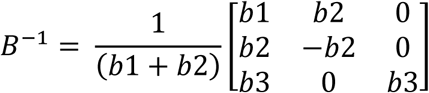

#### Verification: Verification that B × B⁻¹ = I

We verify this by direct matrix multiplication:

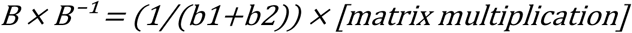

Computing each element of the product matrix:

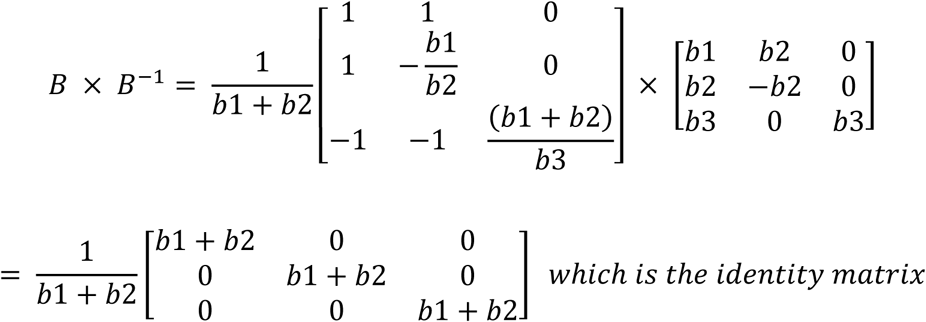

#### Step 2: Identify the mechanism-isolating stimuli

Reading off the columns of B⁻¹, we obtain the three isolating stimuli (LMS vectors that modulate only one DKL mechanism while leaving the others unchanged):

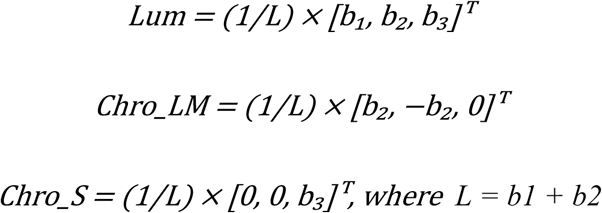

Each stimulus modulates one mechanism without affecting the others. For instance, Chro_LM increases L-cone and decreases M-cone excitation by equal luminance amounts, keeping both the luminance (L+M) and S-cone excitation constant.

#### Step 3: Compute pooled cone contrast for each isolating stimulus

Pooled cone contrast measures the total excursion across all three cone types, normalized by their background excitations:

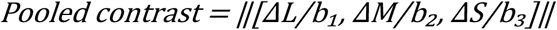

Applying this to each isolating stimulus:

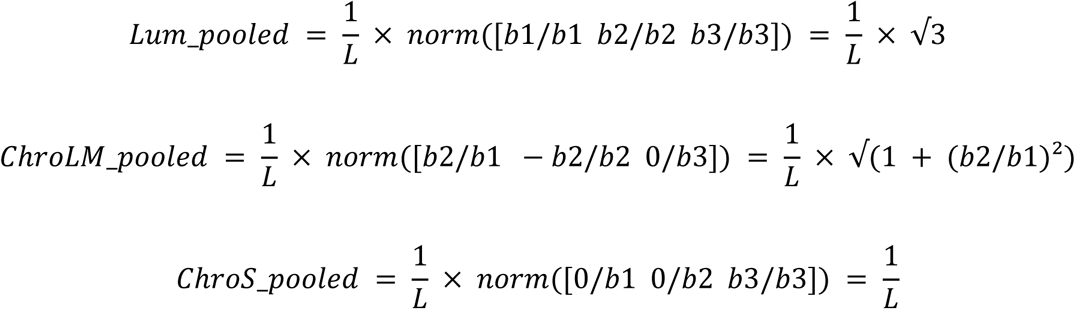

#### Step 4: Normalize to unit pooled contrast

Dividing each isolating stimulus by its pooled contrast yields unit stimuli:

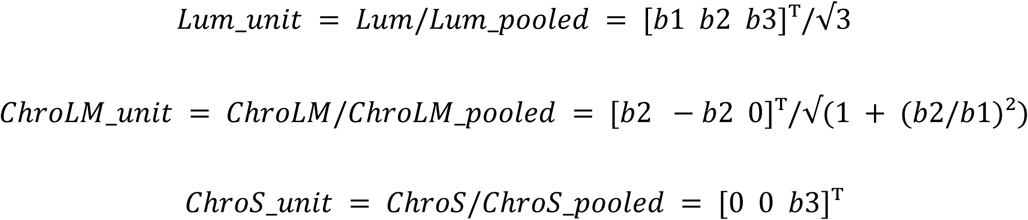

#### Step 5: Computing the normalization constants

Lum_norm = B × Lum_unit:

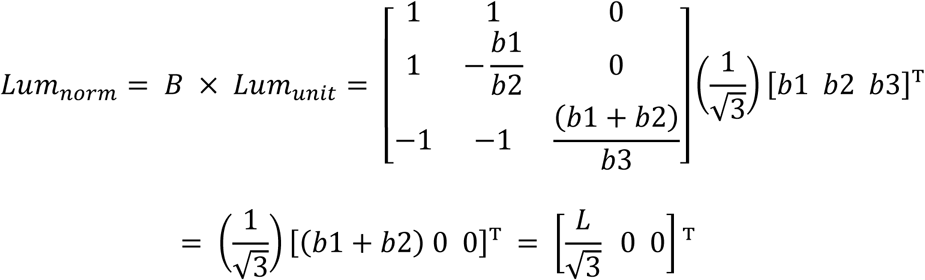

The normalizing constant is the inverse of the first element: √3/L

Similarly, ChroLM_norm = B × ChroLM_unit. Let c = √(1 + (b2/b1)²):

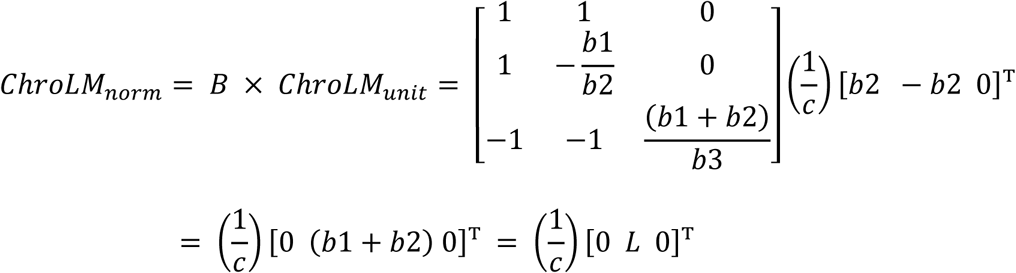

The normalizing constant is the inverse of the second element: c/L = √(1 + (b2/b1)²)/L

Finally, ChroS_norm = B × ChroS_unit:

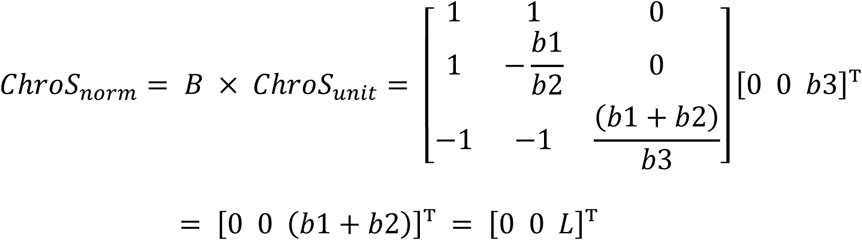

The normalizing constant is the inverse of the third element: 1/L

Hence, the full normalizing matrix D is:

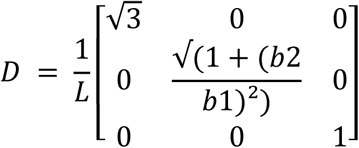

### Implementation

We implemented the complete color transformation pipeline using modified Psychtoolbox (MATLAB) and Lablib (Objective-C) functions, verifying that both approaches produce equivalent results. Key functions include RGBToXYZMatrix (computes the RGB-to-XYZ transformation), XYZToLMSMatrix (provides the XYZ-to-LMS transformation in luminance units), and LMSToDKLMatrix (computes the complete D × B transformation). All calibration code, stimulus generation scripts, and validation data are available at the following Github project repository: https://github.com/supratimray/displayDKLSpace.

